# Phage-plasmid hybrids are found throughout diverse environments and encode niche-specific functional traits

**DOI:** 10.1101/2024.06.18.599647

**Authors:** J. Mullet, L. Zhang, A. Pruden, C.L. Brown

## Abstract

Phage-plasmids are unique mobile genetic elements that function as plasmids and temperate phages. While it has been observed that such elements often encode antibiotic resistance genes and defense system genes, little else is known about other functional traits they encode. Further, no study to date has documented their environmental distribution and prevalence. Here, we performed genome sequence mining of public databases of phages and plasmids utilizing a random forest classifier to identify phage-plasmids. We recovered 5,742 unique phage-plasmid genomes from a remarkable array of disparate environments, including human, animal, plant, fungi, soil, sediment, freshwater, wastewater, and saltwater environments. The resulting genomes were used in a comparative sequence analysis, revealing functional traits/accessory genes associated with specific environments. Host-associated elements contained the most defense systems (including CRISPR and anti-CRISPR systems) as well as antibiotic resistance genes, while other environments, such as freshwater and saltwater systems, tended to encode components of various biosynthetic pathways. Interestingly, we identified genes encoding for certain functional traits, including anti-CRISPR systems and specific antibiotic resistance genes, that were enriched in phage-plasmids relative to both plasmids and phages. Our results highlight that phage-plasmids are found across a wide-array of environments and likely play a role in shaping microbial ecology in a multitude of niches.

**IMPORTANCE:** Phage-plasmids are a novel, hybrid class of mobile genetic element which retain aspects of both phages and plasmids. However, whether phage-plasmids represent merely a rarity or are instead important players in horizontal gene transfer and other important ecological processes has remained a mystery. Here, we document that these hybrids are encountered across a broad range of distinct environments and encode niche-specific functional traits, including the carriage of antibiotic biosynthesis genes and both CRISPR and anti-CRISPR defense systems. These findings highlight phage-plasmids as an important class of mobile genetic element with diverse roles in multiple distinct ecological niches.

## INTRODUCTION

Vehicles of horizontal gene transfer (HGT), such as plasmids and phages, are key drivers of prokaryotic adaptation and evolution (1, 2). In this regard, their role in the mobility of accessory genes, i.e., genes that are not required for the basic life cycle of a mobile genetic element (MGE), is of particular interest (2, 3). MGEs can carry accessory genes encoding diverse traits that may be advantageous to their hosts, including antibiotic resistance genes (ARGs), virulence factors, defense systems such as CRISPR-Cas, metal resistance genes (MRGs), and toxin-antitoxin systems, among many others (3). Such genes can provide hosts with resiliency in the face of changing selective pressures. While MGEs are typically categorized as independent classes (2,4), there is an emerging awareness of inter-element conflicts that can occur between MGEs within individual bacterial hosts (5,6,7,8). For example, some phages, plasmids, and integrative and conjugative elements carry genes encoding defense systems that interfere with the function of co-infecting MGEs (9). Prokaryotic defense systems like these are hypothesized to be acquired through selective bacteriophage predation and have been demonstrated to cluster with and potentially increase the spread of ARGs (10, 11). The carriage of defense systems by MGEs can result in complex ecological and evolutionary dynamics within their host and can significantly alter the community dynamics of microbial populations (9, 10).

Phage-plasmids (P-Ps) are a newly characterized hybrid class of MGE that occupy a unique place in the landscape of prokaryotic genomic elements. These elements can be generally described as temperate (i.e., integrated) phages that retain the ability to replicate in a plasmid-like manner as extra-chromosomal DNA as part of their host life cycle (12). A small set of P-Ps have been shown experimentally to employ a unique combinatorial replication strategy, leveraging both phage lysis and reinfection and the multi-copy number potential of plasmids (12,14). Additionally, P-Ps have been shown to transfer ARGs, certain defense systems, and additional accessory genes from both phages and plasmids (13, 14, 15). With supporting research indicating that P-Ps are significant promoters of genetic exchange between phages and plasmids, the composition and diversity of their accessory genomes remains a key knowledge gap (15). Their unique biology makes the question of their accessory genome particularly intriguing, with the potential for distinct infection and HGT strategies. P-Ps thus represent a distinct class of MGE, a poorly understood dimension of microbial community dynamics, and, hypothetically, a new mechanism of transfer for accessory genes such as AMR or CRISPR-Cas systems.

However, to date, the environmental distribution of P-Ps has not been determined. Indeed, whether P-Ps are common features of microbial communities or merely rare oddities that emerge in specific niches has not yet been ascertained. This limited examination into P-P biological diversity becomes critical to understand as phages and plasmids independently possess unique functional variation across different environments (16, 17). Understanding the diversity of P-Ps across these environments can provide improved insights into the impacts and potential interactions these elements have in the genetic exchange of accessory genes between phages and plasmids.

Here, we recovered a unique dataset of 5,742 P-Ps across four public databases of plasmids and phages and found that they are remarkably prolific across a diverse array of environments. Examination of P-P accessory genome contents suggests a strong linkage to niche-specific ecological dynamics. Compilation and annotation of P-P genomes herein expands knowledge of their genomic diversity and provides new insight into their unique biological and ecological function.

## RESULTS

### P-P hybrids are prolific in public databases of phages and plasmids

We analyzed 1,179,858 genomes from databases of plasmids and phages (PLSDB (19), GPD (20), MGV (21), and IMG/VR (22)) for putative P-Ps using a random forest classifier. The features of the model included the number of hallmark protein hits to each class of MGE (bacteriophage, plasmid, integrative elements, insertion sequences, and multiple), the associated mobileOG-db major categories for each protein, and the number of total proteins and open reading frames for each genome (Supplementary Methods; Table S1) and were trained on (10,289 genomes from [Pfiefer et al.])(12, 18). The final classifier demonstrated an accuracy/FPR/FNR of 95.4%, 2.9%, 19.7%, respectively and was found to outperform manual assignment based on proportions of phage and plasmid-associated gene content (Supplementary Methods; Table S1). This model was employed to generate a conservative, high-confidence set of P-Ps, which was especially relevant because of our usage of IMG/VR v4.0, a database of phage genomes derived primarily from environmental metagenomes (22).

The final P-P dataset examined in this study was composed of 5,742 dereplicated genomes with 137 from GPD, 13 from MGV, 4,425 from IMG/VR, and 1,167 from PLSDB (Fig 2A) (19, 20, 21, 22). PLSDB was predicted to contain several phage genomes (0.8% of PLSDB sequences) and phage databases, such as IMG/VR, were found to harbor many plasmid sequences (Fig 2A) (19, 22). This is not necessarily surprising, as accuracy of plasmid and phage identification can be affected by both low-quality annotated databases and the inherent bias of tools and datasets that specifically classify only one type of MGE. Prior studies have shown that plasmid classification tools can be prone to misidentifying phages as plasmids and, likewise, phage identification tools sometimes misidentify plasmids as phages (18, 28). These inherent biases of analyses targeting a single class of MGE highlight the value of predicting multiple MGE classes simultaneously.

**Figure 1.**
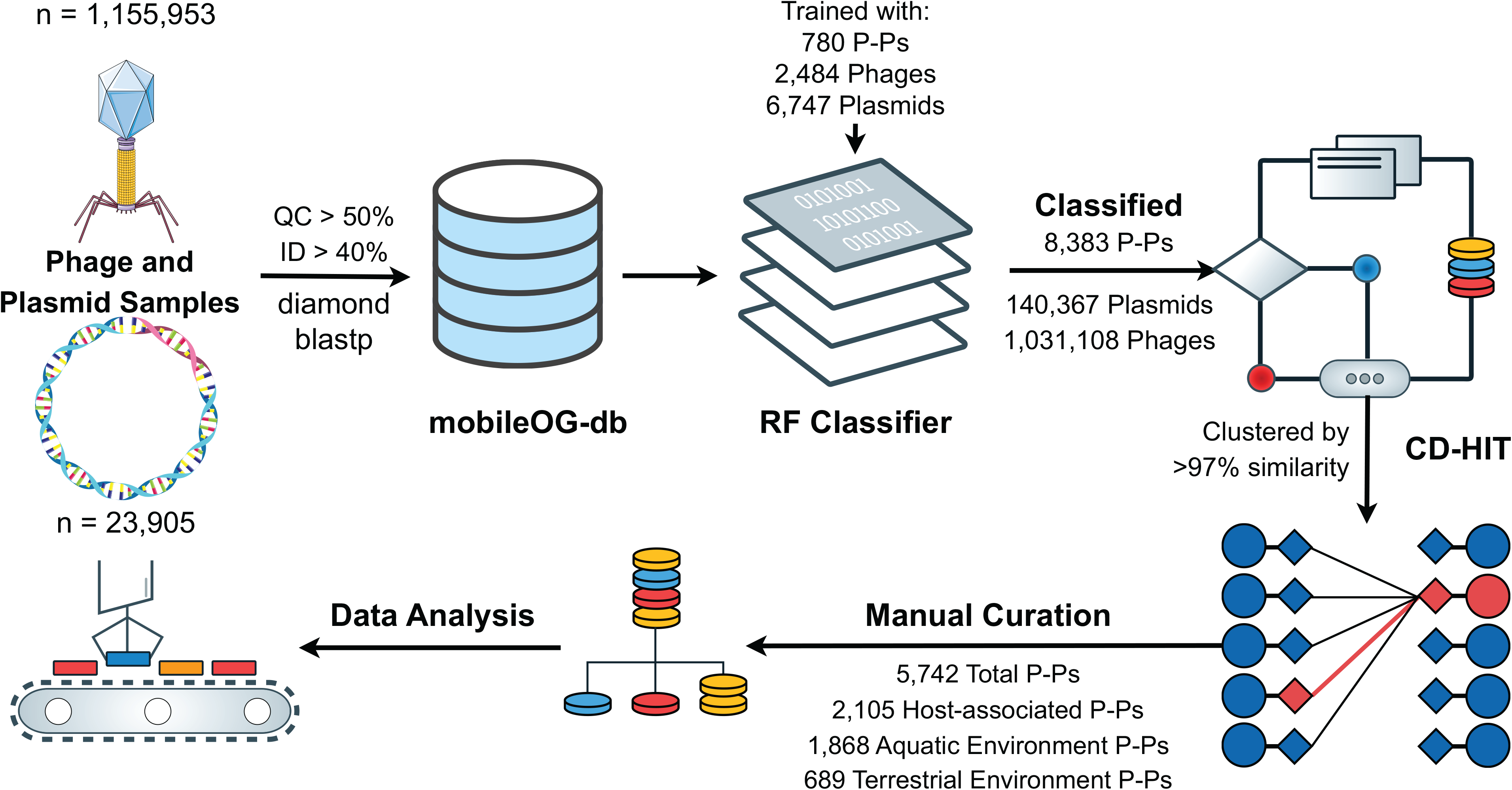
Filtering and identification of phage-plasmids from publicly-available phage and plasmid genomes. The genomes from the three phage databases (n=1,155,953) and one plasmid database (n=23,905) were processed against mobileOG-db to identify MGE-related hallmark genes (18). The genomes were then reclassified into phages (n=1,031,108), plasmid (n=140,367), and phage-plasmid (n=8,383) using a random forest classifier that identifies P-Ps using phage and plasmid hallmark proteins. The phage plasmids were then clustered (n=5,742) to remove identical genomes and manually curated by the associated source location of the classified genomes.

**Figure 2.**
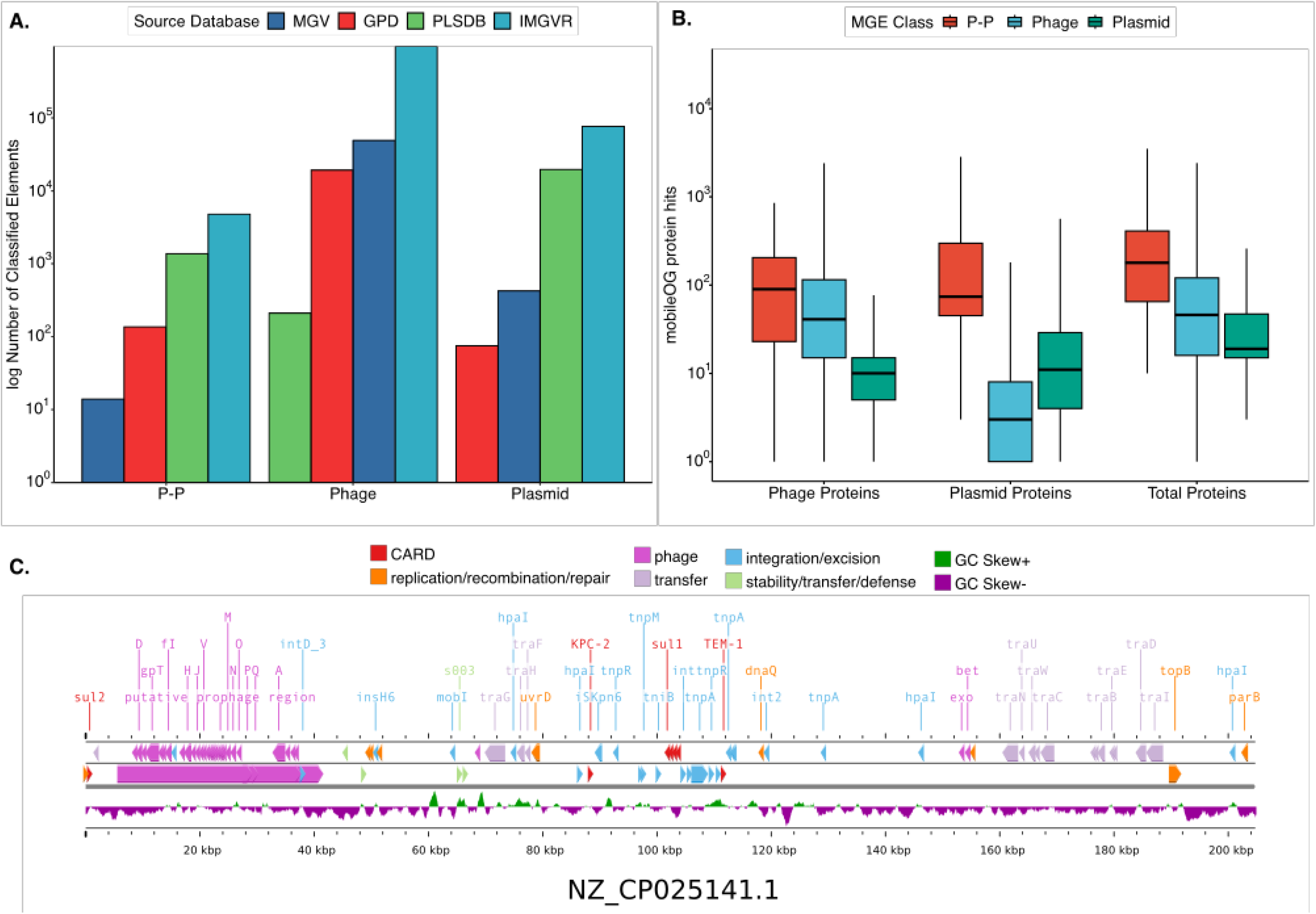
Phage-plasmids (P-Ps) are prolific in databases of plasmids and phages. (A) Number of classified MGEs of each element class from the four respective databases before dereplication. (B) The hybrid nature of P-Ps are reflected in the patterns of mobileOGs. (C) Illustration of a phage-plasmid from PLSDB (id=NZ_CP025141.1) depicted using Proksee including Phigaro, Prokka, mobileOG-db, CARD, and GC Skew annotations (18, 19, 23, 24, 25, 26, 27). All unlabeled or unclassified proteins were removed from this figure.

Metadata across the P-P set was harmonized to group P-Ps according to the environment from which the original sample was sourced: terrestrial (n = 689); aquatic (n = 1,868); host-associated (n = 2,105); and unclassified (n = 1,080) (Supplementary Methods; Table S1). Comparative analysis of mobileOGs (i.e., MGE hallmark genes) highlighted distinct profiles of gene content across phages, plasmids, and P-Ps (Fig. 2B). These profiles were consistent with expectations in that P-Ps encoded more phage genes than plasmids (median 88 genes vs. 8 genes; p < 0.001); more plasmid genes than phages (55 genes vs. 0 genes; p < 0.001); and more total genes than both phages and plasmids (179 P-P genes vs. 46 phage genes vs 19 plasmid genes; p < 0.001) (Fig. 2B). In addition, P-Ps were found to have larger average genome sizes than either phages or plasmids, as has been reported previously in studies that examined a smaller dataset of P-Ps (Supplementary Methods; Fig. S3, 12).

### P-Ps are associated with disparate hosts and ecological niches

Examining the putative hosts of P-Ps can provide insight into the ecology of P-Ps across distinct environmental niches. A compilation of source database metadata was used in tandem with sequence analysis to identify predicted host taxonomy, plasmid incompatibility groups, phage morphology, and the source environment of the P-P genomes. Only 820 (14.3%) genomes were able to be placed within archived plasmid incompatibility groups, likely due to underrepresentation of incompatibility groups beyond *Enterobacteriaceae* in reference databases (30) (Supplementary Methods; Table S1). A putative viral taxonomic classification was obtained for 5,182 genomes classified into viral taxonomic families using geNOMAD (19). The bacterial host taxonomy was obtained with 58.8% of P-Ps (n=3,371) receiving a phylum-level classification (Supplementary Methods; Table S1).

We next investigated the prokaryotic hosts of P-Ps across different environments. The most commonly predicted bacterial host phyla across all environments were *Pseudomonadota* and Firmicutes. The aquatic P-Ps possessed the highest diversity in predicted prokaryotic host phyla, including several phyla (*Verrucomicrobia*, *Crenarchaeota*, and *Euryarchaeota*) exclusively associated with aquatic P-Ps (Fig 3). Further examination revealed differences in the class-level taxonomy of the P-P bacteria. Within the *Pseudomonadota* phylum, *Gammaproteobacteria* was the most common predicted bacteria class, particularly in host-associated P-Ps (95.1% host-associated P-Ps, 76.3% terrestrial P-Ps, and 56.5% of aquatic P-Ps). *Alphaproteobacteria* and *Betaproteobacteria* classes were associated with more terrestrial and aquatic P-Ps (4.8% host-associated P-Ps, 23.6% terrestrial P-Ps, and 46.2% of aquatic P-Ps). The terrestrial P-Ps in *Gammaproteobacteria* were primarily from the *Pseudomonadales* order, while aquatic P-Ps were affiliated with a broader array of bacterial carriers (Fig 3). Host-associated P-Ps were predominately carried by *Enterobacteriaceae* (71.2% of *Pseudomonadota* -associated hosts), a family that includes many enteric Gram negatives of clinical relevance, such as *Escherichia*, *Salmonella*, and *Shigella* (Fig. 3)(31). The *Enterobacteriaceae* bearing P-Ps were more frequently found among the host-associated P-Ps compared to both the aquatic (22.9%) and terrestrial (25.0%) *Pseudomonadota* bearing P-Ps (Fig 3).

**Figure 3.**
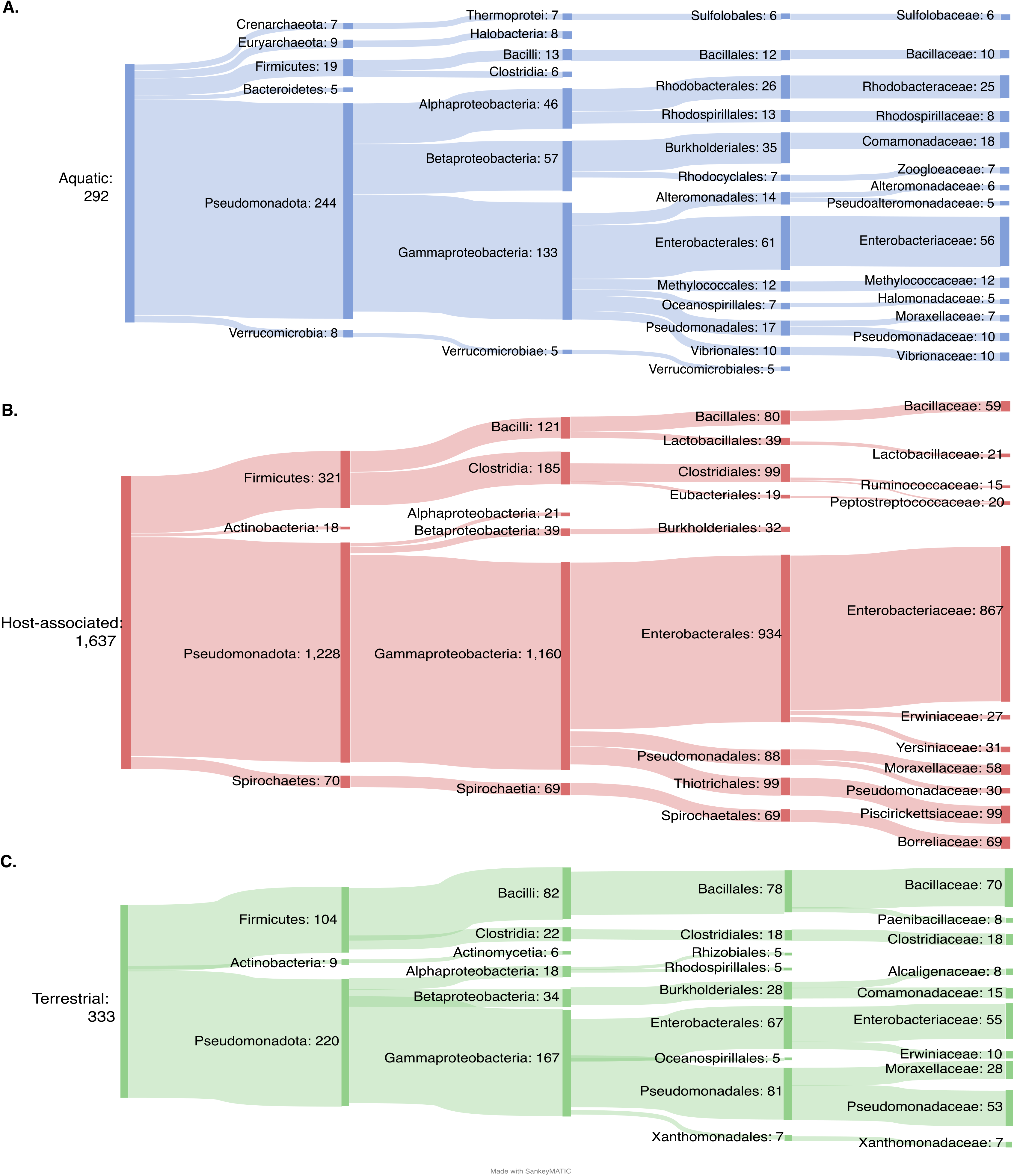
Classification and distribution of P-Ps according to the reported source organism and environment. Examination of the relative abundance of P-P host predicted taxa for aquatic (A), host-associated (B), and terrestrial (C) genomes. The predicted host phyla, class, order, and family of each respective source location are included in each subfigure (29). The predicted host taxa for any P-Ps without reported environmental source locations were excluded from this analysis. Any infrequent taxa that were <1% abundance in the respective environmental location were not included in this figure.

We next examined taxonomy of the P-Ps themselves. An analysis of the updated ICTV family classifications and plasmid incompatibility groups was performed using geNOMAD and PlasmidFinder, respectively (28, 30, 32). *Caudoviricetes* represented the dominant viral order across all environments, with 1.6% of aquatic P-Ps assigned classifications from *Megaviricetes* – an order containing giant viruses (33). At the family-level, few P-Ps could be classified using geNOMAD, but it was noted that the most frequently detected viral family was *Kyanoviridae* (n=70), which was only found among aquatic P-Ps (28). The plasmid incompatibility groups were similar across different environments, with IncFIB, IncY, and p0111 being the most common classifications.

### P-Ps encode diverse and niche-specific accessory functions

The broad distribution of P-Ps across disparate environments led us to question what functional traits P-Ps might carry across a correspondingly wide variety of ecological niches. We next investigated the accessory genome of P-Ps, including ARGs, metabolism-related genes, metal resistance genes, defense systems, toxin-antitoxin systems, anti-CRISPR systems, and virulence factors.

Accessory gene content of P-Ps was relatively unchanged within each of the distinct environments from which the P-Ps originated (Fig. 4). When comparing P-Ps to phage and plasmid accessory genes, P-P accessory gene profiles were most similar to those of plasmids (Kruskal-Wallis and post hoc Dunn test; p=5.30 x 10^-1^), with very few accessory genes found among phages relative to plasmids and P-Ps (Kruskal-Wallis and post hoc Dunn test; p= 1.15x 10^-9^) (Fig. 4). However, it was noted that the P-Ps had enriched anti-CRISPR genes compared to phages and plasmids (Fischer exact test; 240 P-P genes vs. 5 phage genes vs 1 plasmid genes; p < 0.001). While most ARGs, MRGs, and virulence factors likely predominately originated from plasmid sources, it is also possible that phages still contribute to certain metabolism and defense system accessory genes among P-Ps.

**Figure 4.**
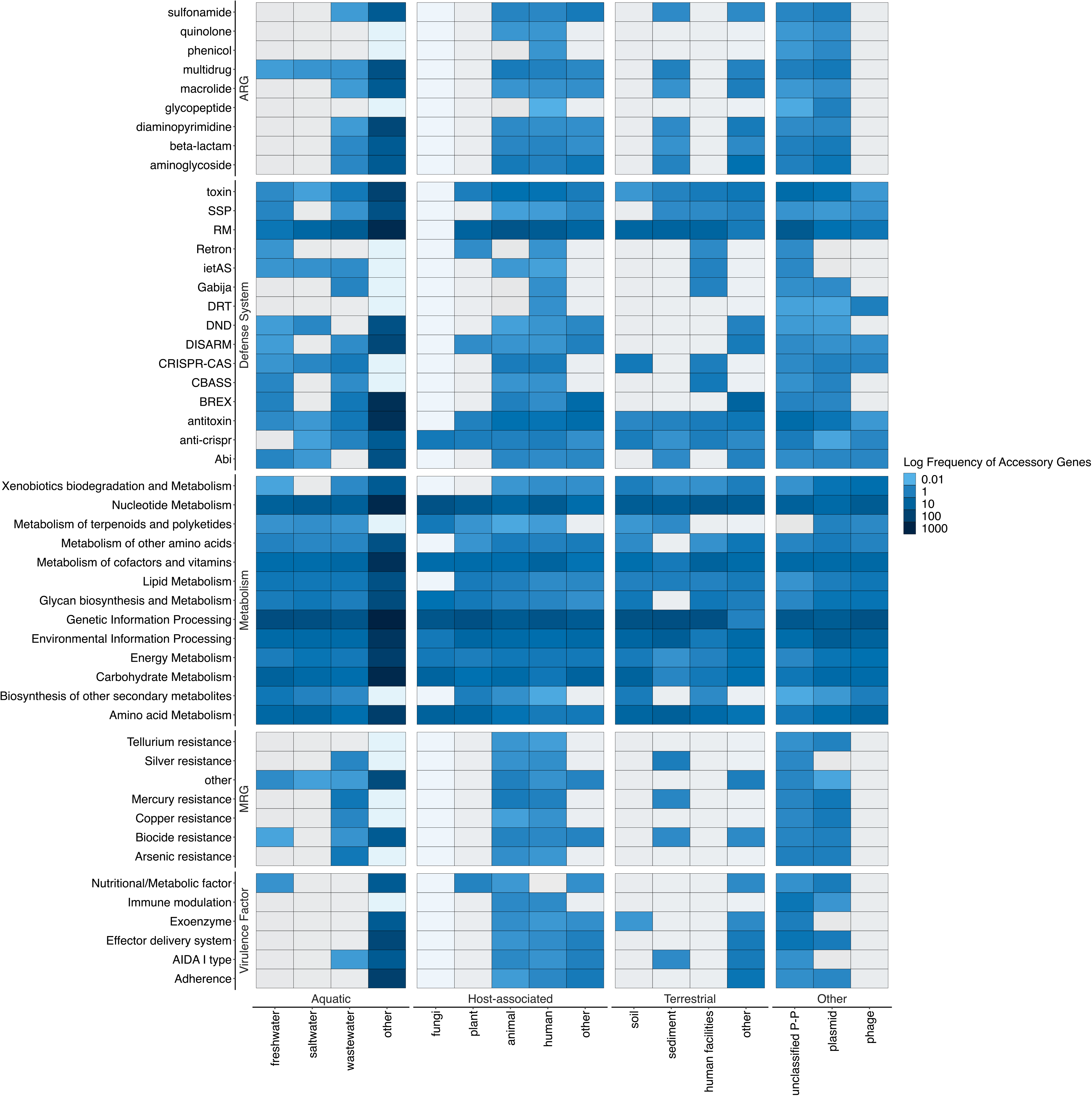
Comparative analysis of key accessory genes found to be carried by the P-Ps across a diverse range of source environments. The accessory genes were grouped into virulence factors, metal resistance, metabolism, defense systems, and antibiotic-resistance genes (ARGs). These genes were grouped into associated functional categories as shown in the supplementary tables. It was noted that both the toxin-antitoxin genes identified from TADB and the anti-CRISPR genes classified from Anti-CRISPRdb v2.2 were grouped with the defense system genes for visual purposes (34, 35). Only accessory gene categories with at least 25 hits were included in the figure above. The values were taken from the log10 of the relative frequency of the genes compared to the total number of accessory genes found in each element source location. The plasmid and phage categories comprise 500 random phages and plasmids, capturing differences between the various class of MGEs and acting as experimental baseline controls for comparing phages, plasmids, and P-Ps.

### Diversity within the unique accessory genomes of phage-plasmids

We sought to further characterize the diversity among the unique P-P accessory genomes and to assess additional differentiating features and trends among their profiles. First, we analyzed the differences between P-P and plasmid ARG gene distributions. Similar to prior research, it was noted that P-Ps possess ARGs less frequently than plasmids. However, some ARGs, including cpxA, EcoI_emrE, and CTX-M-142, were enriched in P-Ps compared to plasmids (Fischer exact test; p < 0.01) (Supplementary Methods; Fig. S6). We found that several of the most common ARGs are associated with Class I integrons, including sul1, aadA2, and qacEdelta1 (Fig. 5a). While approximately 5% of the host-associated P-Ps contained ARGs, the aquatic and terrestrial environment phage-plasmids appeared to be more depleted in the number of ARGs (38). It was noted that the P-Ps associated with wastewater environments contained a few ARGs possessing the *blaCTX-M-15* gene, which is one of the most common extended-spectrum beta-lactamase (ESBL)-encoding ARGs found to be associated with infections that are resistant to third-generation cephalosporins (37). Through the visualization of genetic contexts surrounding CTX-M-15, we found a conserved region that was encountered in P-Ps encountered across several examined source environments (Fig. 5c).

**Figure 5.**
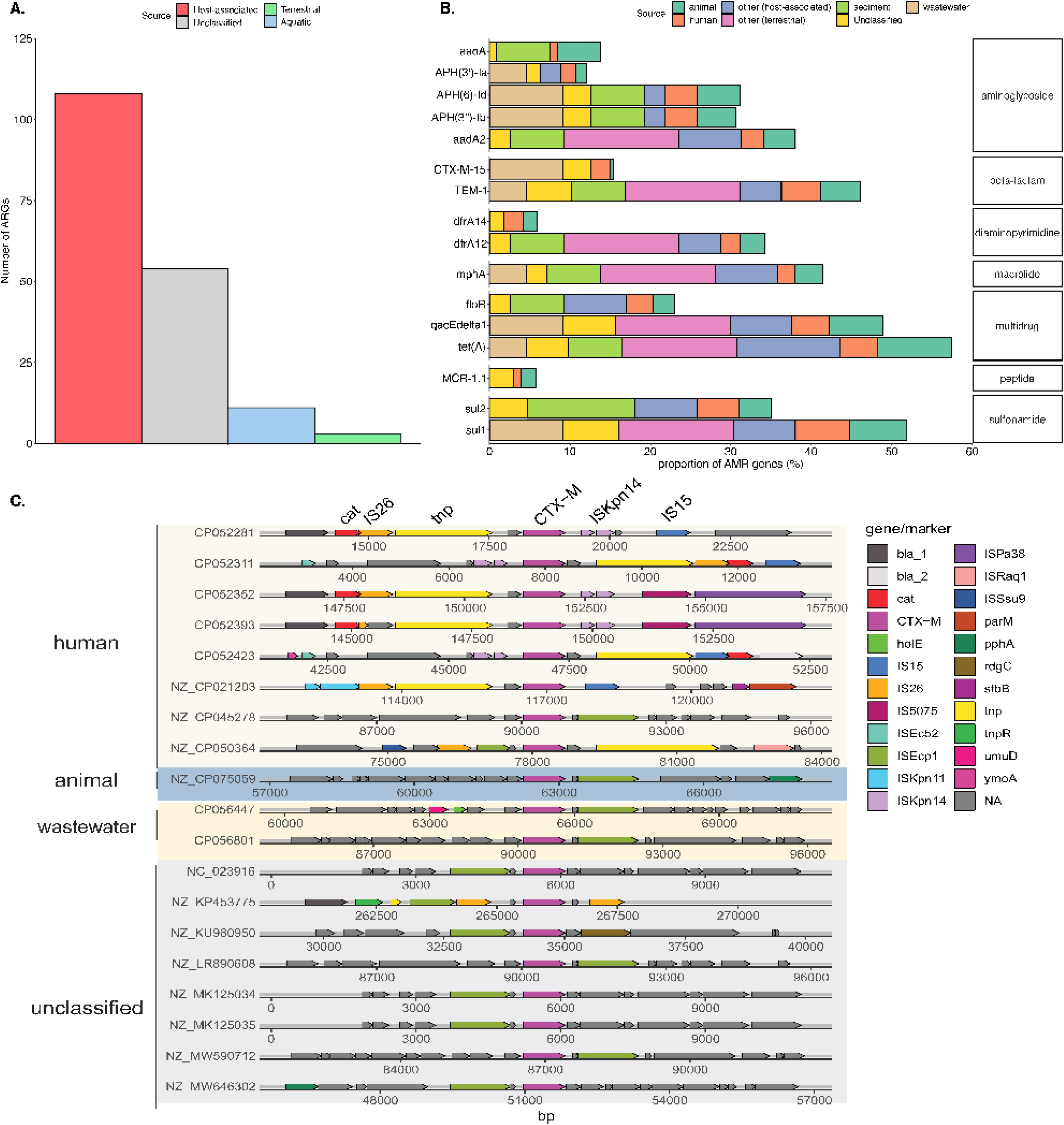
Diversity and distribution of ARGs among P-Ps of various origin. (A) Total number of all ARGs found in P-Ps originating from each source environment. (B) Frequency of common antibiotic resistance genes (ARGs) carried by the P-Ps relative to the total identified ARGs in each source environment. Only source environments possessing >8 unique ARGs were included in the figure. (C) Gene-to-gene alignment of the CTX-M-15 ARG grouped by the respective source environment of the phage-plasmids (36).

Because of the hybrid status of P-Ps as having both phage and plasmid type genes, an intriguing question is whether they utilize distinct defensive and offensive systems for interelement competition. We assessed the diversity of defense system genes including both CRISPR and anti-CRISPR systems. From this examination, P-Ps were found to possess more anti-CRISPR systems compared to CRISPR-Cas systems (Fig. 6a) across all environments. We determined that only one P-P (NZ_CP063966.1) possessed both a CRISPR-Cas system and anti-CRISPR system (Supplementary Methods; Fig S7). This indicates that P-Ps typically utilize only one of these defensive or offensive strategies for limiting additional MGE co-infection (39, 40). The reduced variation of both systems in P-Ps was also noted. CRISPR-Cas systems genes were only found in Class I (n=132), Class III (n=32), and Class IV (n=36) among the five major categories. Interestingly, the predominant anti-CRISPR system genes detected was the AcrIIA7 (n=233), one of the most abundant anti-defenses CRISPR-associated inhibitors (41).

**Figure 6.**
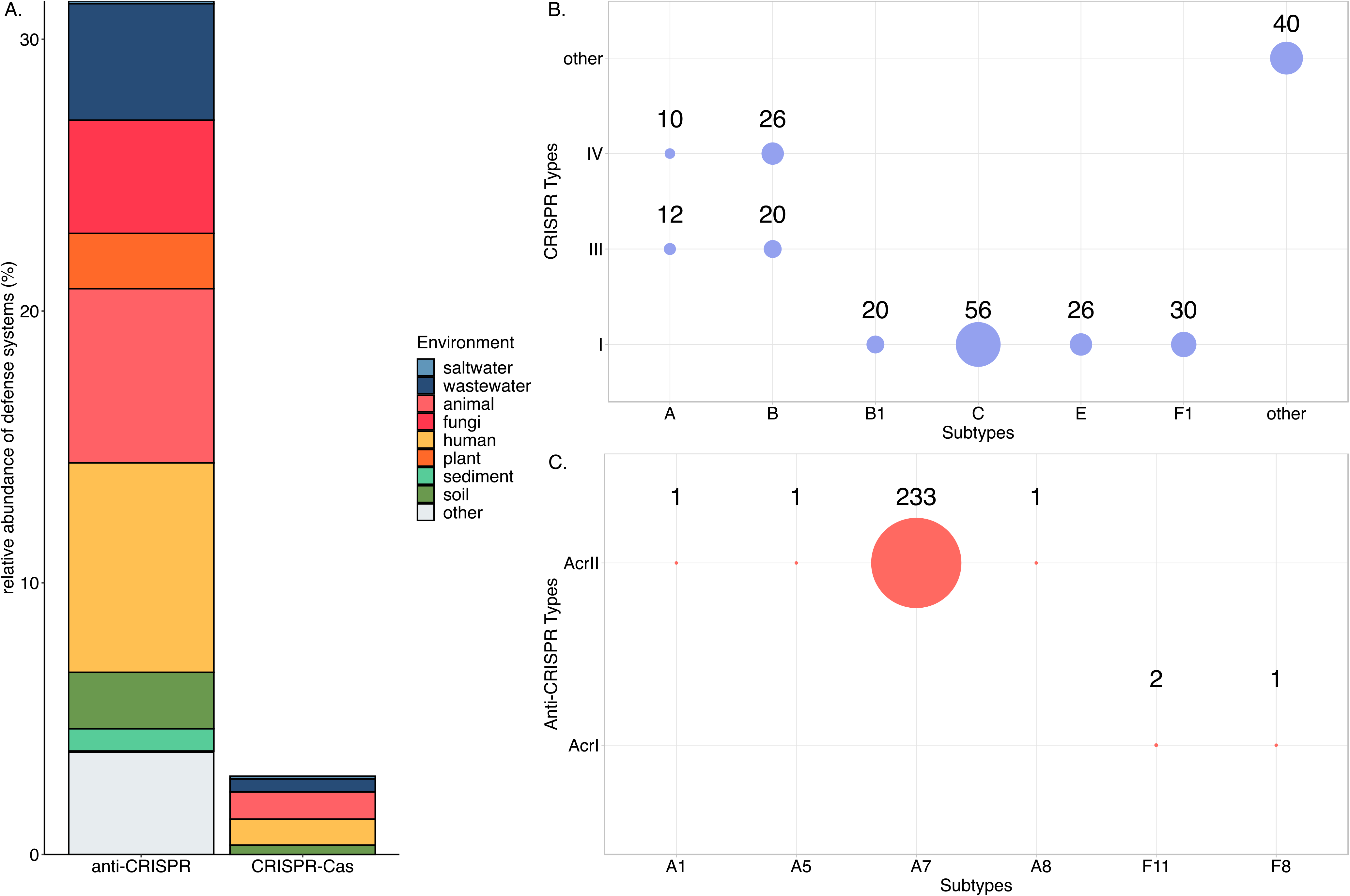
Analysis of the diversity and frequency of CRISPR-Cas and anti-CRISPR systems. (A) Relative abundance of anti-CRISPR and CRISPR-Cas systems encountered among unique P-P genomes reconstructed from each of the respective source environments (Relative Abundance = P-P genomes containing defense system from a respective source location / Total P-P genomes from respective source environments). (B) Distribution and occurrence of total CRISPR-Cas gene subtypes in P-Ps. “Other” includes systems that could not be classified into one single category or which were classified as a category other than the five primary classes of CRISPR-Cas systems. (C) Prevalence and abundance of anti-CRISPR system genes in P-Ps. Only subtypes found in the P-Ps are displayed in the figure.

CRISPR-Cas defense systems were frequently found in host-associated and terrestrial P-Ps, with lower abundance among the aquatic P-Ps (Fig. 6). Most environments were characterized by even abundance of both classes of defense systems. However, some samples only recorded examples of one defense system class, such as animal host-associated P-Ps that possessed only CRISPR-Cas systems and fungi and plant P-P genomes that carried anti-CRISPR systems. These results demonstrate that P-Ps can carry CRISPR-Cas and anti-CRISPR systems in various environmental sources, however, these defense systems appear to be most commonly encountered in host-associated P-Ps.

After examining the unique contributions of ARGs, CRISPR-Cas systems, and anti-CRIPSR systems to phage-plasmid accessory genes, it appeared that many of these genes are relatively consistent in their distribution across all environments from which the P-Ps were derived. However, ARGs and certain defense systems were more abundant among host-associated P-Ps. The metabolic accessory genes were then examined to further investigate how this trend could impact other accessory gene functions. It was noted that the host-associated P-Ps possessed higher abundances of ARGs, CRISPR-Cas systems, anti-CRISPR systems and specific metabolic pathways such as pyrimidine metabolism, drug resistance, and cofactor and vitamin biosynthesis (Fischer Exact Test with a Benjamini-Hochberg correction; p < 0.001) (Supplementary Methods; Fig. S9). The freshwater and saltwater P-Ps contained enriched macrolide biosynthesis, photosynthetic genes and unique nucleotide metabolic pathways such as polyketide sugar biosynthetic pathways (Fischer Exact Test with a Benjamini-Hochberg correction; p < 0.001) (Supplementary Methods; Fig. S9).

To further examine the diversity of the metabolic accessory genes found on P-Ps, we considered the dTDP-6-deoxy-α-D-allose biosynthesis pathway. This pathway is a critical for the formation of mycinose, as dTDP-6-deoxy-α-D-allose is the last free intermediate in this biosynthesis pathway (46). Mycinose is an important biomolecule that assists in forming several macrolide antibiotics (46). The P-Ps noted to possess this metabolic pathway were exclusively aquatic P-Ps and they all contained identical KEGG Modules (M00794) (45). In particular, these aquatic P-Ps contained three of the four enzymes in this pathway, including dTDP glucose 4,6-dehydratase, an enzyme that assists in forming all 6-deoxy sugar biosynthesis (Fig. 7) (47). These P-Ps contained genes encoding two enzymes (dTDP-4-dehydro-6-deoxy-D-glucose-3-epimerase and dTDP-4-dehydro-6-deoxy-α-D-gulose-4-ketoreducatase) that are essential to the dTDP-6-deoxy-α-D-allose biosynthesis pathway (42). The presence of the intermediate steps of the nucleotide sugar pathways (e.g., Fig. 7) suggests that P-Ps could stimulate auxiliary metabolite production from host-derived inputs of glucose 1-phosphate, dTTPs, and thymidyltransferase. Many polyketide sugars are frequently associated precursors for bacterial-produced antibiotic pathways, and these were exclusively found in aquatic P-Ps.

**Figure 7.**
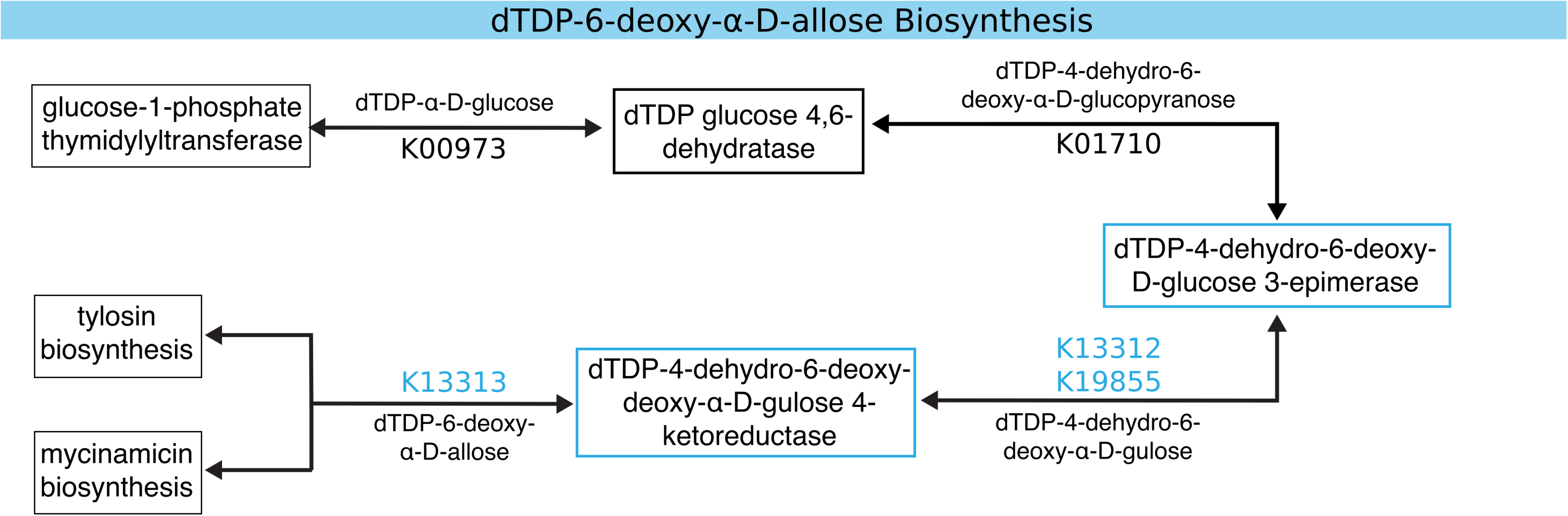
dTDP-6-deoxy-α-D-allose biosynthesis pathway found in some aquatic P-Ps (n=14/1,868) (**42, 43, 44**). The blue-outlined boxes indicate the portion of the associated pathway found in the P-Ps. Among the 14 P-Ps found to carry portions of this pathway, 13 were derived from freshwater and one from saltwater. The designated KEGG pathways align with the reaction products from these enzymes with the blue KEGG pathways indicating portions of the pathway that the P-P carried accessory genes (45).

## DISCUSSION

Here, we investigated the functional repertoire of accessory genes and the ecological diversity of P-Ps. P-Ps were found to inhabit a wide range of environments and exhibited notable genetic variation, with evidence suggesting that most accessory genes are derived from plasmids (12, 13, 15). P-P encoded accessory genes included a diverse arsenal of ARGs, CRISPR-Cas systems, virulence factors, and metabolism genes. While prior research primarily demonstrates that P-Ps possess most accessory genes at rates intermediate to both phages and plasmids, we found evidence that some accessory gene elements are disproportionately associated with P-Ps (12, 13, 15). Specifically, we found that anti-CRISPR systems and some ARGs [cpxA, EcoI_emrE, and CTX-M-142] were enriched in these elements (Supplementary Methods; Fig. S6, Fig. 6). With our developing understanding of MGE competition (e.g., plasmids containing CRISPR-Cas systems that may target bacteriophages), it raises questions about the role of P-Ps in such interactions (9, 48). Prior work has shown that some phages bearing anti-CRISPR systems have density-dependent protection from CRISPR-Cas, suggesting a role for cooperation and/or co-infection in the defense mechanism (49). Furthermore, it has been observed that P-Ps can exploit the replication machinery of plasmids to achieve a plasmids’ relatively high copy number potential (14, 49). This replication strategy could allow for higher phage densities, thus potentiating anti-CRISPR systems.

P-P accessory gene content differed across environments. We examined various functional genes to investigate whether P-Ps confer traits that assist their prokaryotic hosts in adapting to their local environments. While these are not an exhaustive list of potential accessory genes, they are among the most important in understanding the ecology of P-Ps and their relevance to human health. We found that the distributions of these accessory genes varied significantly across environments. Host-associated P-Ps were enriched with defense systems and ARGs compared to aquatic P-Ps with increased abundances of intermediate secondary metabolic pathway genes. Many of these accessory genes appeared to be conserved, but the frequency depended on the environments from which these elements were sourced (e.g., Fig. 5,6,7). The overall trends of accessory genes appear similar to prior studies investigating plasmid gene diversity, although future works should investigate the differences between plasmid and phage-plasmid accessory genes (38, 50). The variability in accessory gene content among P-Ps suggests that these elements might occupy unique niches within microbial communities depending on their environments.

P-P genomic variation has the potential to alter microbial communities. Through the diversity of accessory gene content in host-associated, aquatic, and terrestrial-sourced P-Ps, we found a wide array of biologically-relevant accessory genes. These elements are prone to recombination and genetic exchange with other MGEs, making them of particular interest when considering their accessory genomes (15). These unique biological features with the diverse array of accessory genes highlight the importance of further study into these elements (12, 13, 14, 15, 50). Our results suggest that P-Ps offer notable genetic diversity and complexity that may impact MGE and bacterial evolution. The inherent variability of their hosts, viral genes, plasmid components, and functional genes these elements possess can play a significant role in shaping the recombination and HGT events in microbial populations. Understanding and potentially monitoring P-P populations offers potential benefits to mechanistic understanding of the recombination and transmission of accessory genes such as ARGs, MRGs, and virulence factors, contributing to their overall spread. The P-P accessory genome should be studied further to fully understand how these elements spread this diverse assortment of accessory genes.

## METHODS

### Data Acquisition and Processing

The complete genomes of 33,595 plasmids were retrieved from PLSDB, 19,510 genomes from GPD, 52,958 genomes from MGV, and 1,416,547 genome and associated fragments from IMG/VR databases (19, 20, 21, 22). An additional 8,248 plasmids, 2,256 phages, and 780 P-Ps were obtained from Pfeifer et al. for training the random forest classifier (12). We removed genomes smaller than 10 kb to remove potentially fragmented genomes and genomes larger than 300 kb to avoid megaplasmids and chromatids. The information regarding the appropriate virus taxonomy, sampling source location, and additional information was collected from the metadata from PLSDB, GPD, MGV, and IMG/VR sources (19, 20, 21, 22). All analyses were conducted in Python (https://www.python.org/) unless otherwise stated.

### Annotation of Protein Sequences

The genomes from PLSDB, GPD, MGV, IMG/VR, and Pfeifer et al. were processed with Prodigal (v2.6.3) using the (-p) meta setting to generate open reading frames (12, 19, 20, 21, 22, 51). The open reading frames were aligned to predicted protein sequences using diamond blastp (v4.6.8) using a minimum identity of 40%, minimum query coverage of 50%, maximum e-score of 1 x 10^-5^, and k value of 15 (51). The less stringent settings allowed for the acquisition of more diverse phage species to ensure a high identification of all MGEs using mobileOG-db (Beatrix v1.6) (18). This database provides an inclusive and diverse distribution of MGE protein sequences, which allows for a robust analysis of MGEs.

### Identification of Phage-Plasmids (P-Ps)

A random forest classifier was trained using the outputted results from the protein alignments using mobileOG-db (18). The features utilized in the classifier included the number of protein hits to bacteriophages, integrative elements, insertion sequences, plasmids, and multiple MGE class proteins. In addition, the associated mobileOG-db major categories of the proteins (phage, integration/excision, replication/recombination/repair, transfer, and stability/transfer/defense) were included with the total number of proteins and ORFs found in each genome (18). The Pfeifer et al. paper used several classification techniques including identifying P-Ps from literature sources, plasmid HMMs found in phages, and plasmids with identified phage-specific profiles for classifying P-Ps due to the limited known P-Ps prior to their work (12). This paper utilizes the prior data obtained to train this classifier with the now known quantity of P-Ps. The model’s training began by performing ten randomized training sets using approximately 20% of samples as test data and 80% as training data. The random forest classifier had a max decision depth of 8 and used entropy as the criteria measurement. The performance results from the ten randomized trials were averaged to examine the effectiveness of the random forest classifier. The classifier achieved an average accuracy of 95.4% and a false positive rate of 2.9%. The testing data consisted of approximately 160 P-Ps, 500 phages, and 1350 plasmids, while the training data contained 620 P-Ps, 2,000 phages, and 5,400 plasmids (12). The PLSDB, MGV, GPD, and IMG/VR genomes were then classified using the trained random forest classifier to identify whether each element was a plasmid, phage-plasmid, or bacteriophage (19, 20, 21, 22). CD-HIT-EST v4.6.8 was utilized to cluster the sequences and remove sequences with < 97% sequence similarity (53). The P-Ps were then examined using CompareM to compare the average nucleotide identity between the samples to compare the sequence similarity after clustering (54).

### Manual Curation of Source Location

The classified phage-plasmid genomes were cross-referenced with the source database metadata to determine additional information regarding the source locations and taxonomy for additional analysis. The P-Ps were then categorized by environmental source location into the following categories: aquatic, terrestrial, host-associated, and unclassified environments. These categories were separated into more unique categories according to the exact location of the genomes, including the subcategories of saltwater, freshwater, wastewater, other aquatic genomes, soil, sediment, human facilities, other terrestrial, human, fungi, animal, plant, other host-associated genomes, and unclassified genomes. Genomes designated as others had designated source locations but were too generalized to classify the genomes further correctly. Phage-plasmids with undocumented source locations were cross-referenced with NCBI BioSample to classify the elements further, but genomes that still could not be classified were designated as other. All genomes with no metadata source locations or metadata with ambiguous locations were removed from source location analysis.

### Data Analysis

The taxonomy of the P-Ps was classified using the associated source metadata from the respective databases. Due to the limited phage and plasmid taxonomy, the associated incompatibility groups of the P-Ps were further classified using PlasmidFinder (v2.1.6) with default parameters and the viral taxonomic classifications were classified using geNOMAD (v1.5.2) (19, 30). The study further identified the key accessory genes of the phage-plasmids, including ARGs, defense systems, toxin-antitoxin systems, metabolism genes, metal resistance genes, and virulence factors. The defense systems were identified using PADLOC (v.1.1.0) classification tool (55). The phage-plasmid genomes were processed through GhostKoala to extract the KEGGs from the Reconstruction Mapper function for identifying the metabolic genes (45, 56). Microbe Annotator (light-v2.0.5) was used to identify complete or partially complete KEGG Module pathways from the specific P-Ps using the blast settings (45, 57). These pathways were classified if the P-Ps contain 50% of the required genes for a specific biosynthesis pathway.

The virulence factors, metal resistance genes, anti-CRISPR genes, and the toxin-antitoxin systems were classified by processing the phage-plasmids using Diamond blastp (v4.6.8) against the VFDB genes from set A, the BacMet2 Predicted dataset, Anti-CRISPRdb (v2.2) database, and the TADB (v.2.0) database with query coverage of 80%, percent identity of 90%, and e-score of 1×10-5 (34, 35, 52, 58, 59). The classified phage, plasmid, and P-P genomes were queried against CARD (v3.0.7) with a minimum identity of 80% and an e-value<10^-10^ (26). The phage-plasmid genomes were then processed through EggNOG-Mapper (v2) to get the associated PFAMs, and COGs for the additional P-P analysis (60, 61, 62). A random selection of 500 phages and 500 plasmids were isolated from the prior classified phage and plasmids classifications. These genomes were processed utilizing the same tools as the phage-plasmids to determine the accessory genes found in these genomes. The graphical analysis was performed using R (https://www.r-project.org/), draw.io (http://draw.io/), and bioicons (https://bioicons.com/).

## Data Availability

All available data can be download from the databases analyzed in this study with all associated accession identification numbers located in the supplemental tables. The supplementary data can be found at the manuscript FigShare repository located at: https://figshare.com/s/b0ffbc71c0bf43e251df. Scripts used in data mining, processing the data, and generating the scripts can be found at https://github.com/jamesm224/phage-plasmid-classification.

## Acknowledgements

We would like to express our gratitude and appreciate for all of the assistance and help that went into designing this study. The authors acknowledge the Advanced Research Computing at Virginia Tech for providing the computational resources used in this study. This study was supported by NSF CI4-WARS (PI Zhang) Award 2004751, the NSF NRT-HDR training (PI Pruden) Award 2125798, and the Fralin Life Sciences Institute.

The authors report no conflict of interest.

